# Large-scale investigation for antimicrobial activity reveals novel defensive species across the healthy skin microbiome

**DOI:** 10.1101/2024.11.04.621544

**Authors:** Uyen Thy Nguyen, Rauf Salamzade, Shelby Sandstrom, Mary Hannah Swaney, Liz Townsend, Sherrie Y. Wu, J.Z. Alex Cheong, Joseph A. Sardina, Isabelle Ludwikoski, Mackinnley Rybolt, Hanxiao Wan, Caitlin Carlson, Robert Zarnowski, David Andes, Cameron Currie, Lindsay Kalan

**Author notes:** Correspondence: Lindsay Kalan, McMaster University, 1280 Main St. W, Hamilton, ON, L8S 4K1. these authors contributed equally (Uyen Thy Nguyen, Rauf Salamzade, Shelby Sandstrom).

## Abstract

The human skin microbiome constitutes a dynamic barrier that can impede pathogen invasion by producing antimicrobial natural products. Gene clusters encoding for production of secondary metabolites, biosynthetic gene clusters (BGCs), that are enriched in the human skin microbiome relative to other ecological settings, position this niche as a promising source for new natural product mining. Here, we introduce a new human microbiome isolate collection, the EPithelial Isolate Collection (EPIC). It includes a large phylogenetically diverse set of human skin-derived bacterial strains from eight body sites. This skin collection, consisting of 980 strains is larger and more diverse than existing resources, includes hundreds of rare and low-abundance strains, and hundreds of unique BGCs. Using a large-scale co-culture screen to assess 8,756 pairwise interactions between skin-associated bacteria and potential pathogens, we reveal broad antifungal activity by skin microbiome members. Integrating 287 whole isolate genomes and 268 metagenomes from sampling sites demonstrates that while the distribution of BGC types is stable across body sites, specific gene cluster families (GCFs), each predicted to encode for a distinct secondary metabolite, can substantially vary. Sites that are dry or rarely moist harbor the greatest potential for discovery of novel bioactive metabolites. Among our discoveries are four novel bacterial species, three of which exert significant and broad-spectrum antifungal activity. This comprehensive isolate collection advances our understanding of the skin microbiomes biosynthetic capabilities and pathogen-fighting mechanisms, opening new avenues towards antimicrobial drug discovery and microbiome engineering.

## Introduction

Human skin represents a first line of defense against mechanical and chemical insults while maintaining homeostasis^1–7^. One of the most significant roles of the skin is to act as a barrier to invading pathogens. This physical barrier is fortified by a diverse microbiome composed of bacteria, fungi, viruses, and microeukaryotes^8^. The surface area of the skin is estimated to be around 30 m^2^, including appendages such as hair follicles and sweat ducts^5^, making it one of the most expansive direct host-microbe interfaces in the body. Across the body, unique microenvironments form on the skin with characteristic moisture levels, pH, and lipid content, driving the composition of the associated microbial community^7,9–12^. Far from mere bystanders, members of the skin microbiome help maintain the integrity of the skin barrier through both direct and indirect defense mechanisms^9–11^.

The skin microbiome fulfills diverse additional functional roles including establishment of immune tolerance and sensing pathogens^13–15^. Skin microbiota indirectly compete against potential pathogenic invaders by consuming limited nutrients and acidifying the skin surface^1,16^. Constituents of the microbiome also engage in direct defense of the skin through mechanisms such as production of antimicrobial molecules. Recent efforts to mine the human microbiome for genes encoding bioactive metabolites have revealed a rich biosynthetic potential^17^. Given that bacteria inhabit different niches within the human body and are exposed to different competitors in the environment, we expect that skin microbiota produce metabolites that are relevant to the niche that they colonize. An example is lugdunin, a non-ribosomal thiazolidine cyclic peptide produced by *Staphylococcus lugdunensis* which is commonly found in nasal cavities. Lugdunin is bactericidal against methicillin-resistant *S. aureus* and strains of vancomycin resistant *Enterococcus* but shows no toxicity toward primary human erythrocytes or neutrophils^18^. An additional example is *Cutibacterium acnes,* commonly found at sebaceous body sites, which produces cutimycin, a thiopeptide with anti-staphylococcal activity^19^. Further, numerous human skin coagulase-negative *Staphylococcus* commensal species produce lantibiotics that inhibit *S.* aureus^18,20–23^.

Characterization of antimicrobial products from the skin microbiota has historically focused on easily cultivated, high-abundance organisms, particularly *Staphylococcal* species, targeting the Gram-positive skin pathogen *S. aureus*^18–20,24–27^. Recent studies have shown these abundant skin microbes harbor a wide diversity of biosynthetic gene clusters (BGCs) in their genomes^28–30^. However, the skin microbiome also contains numerous species of low abundance^31^ with unexplored biosynthetic potential. These species likely harbor many more uncharacterized BGCs. Given that currently characterized BGCs come from a limited set of microbial genera, and most BGCs remain uncharacterized^32,33^, the skin microbiome represents a potentially rich, untapped source of antimicrobial compounds. This potential is further amplified by the presence of distinct and unique BGCs, likely to encode for the synthesis of novel bioactive compounds. These facts suggest that most of the chemical diversity encoded with skin-associated microbial genomes remains unknown. The ongoing discovery of new bacterial species on the skin through combined metagenomic assembly and cultivation approaches, exemplified by the recent Skin Microbial Genome Collection (SMGC) identifying 174 new bacterial species, further underscores the richness of this ecosystem^31^. Identifying the yet-unknown metabolites produced in this niche will not only deepen our understanding of skin biology but likely unveil novel therapeutic molecules.

We have developed the The EPithelial Isolate Collection (EPIC), a microbial biorepository of 6,540 bacterial strains isolated from 1.060 mammalian samples. This new collection is derived from epithelial sites such as the skin, oral, and nasal barriers in humans, swine, non-human primates, horses, chickens, goats, donkeys, and cows. Here, we detail the extensive characterization of healthy human skin isolates included in EPIC, along with matched metagenomes from eight distinct human skin body sites. This subset of EPIC includes 980 bacterial isolates, 287 whole genomes, and 268 metagenomes (**Figure 1**). Our collection is much larger and significantly more diverse than previous collections^31^, comprising hundreds of rare and low-abundance strains, many of which are not included in previously reported collections^31,34^. We use a large-scale solid-phase co-culture screen to assess pairwise interactions between skin-associated bacteria and potential pathogens spanning Gram-negative, Grampositive and fungal pathogens. This data reveals broad antifungal activity by members of the skin microbiome. Comparing gene cluster families (GCFs) allowed us to visualize hundreds of GCFs and demonstrate most of them are unique to our collection. Finally, we describe four novel skin-associated bacterial species, unique to this collection, that exert significant antifungal activity. The BGCs of these novel species are distinct from those in related species of the same genera. This rich and diverse isolate collection enables systematic exploration of skin microbiome chemistry, illuminates the breadth of antagonistic interactions within the skin microbiome, and will provide new molecular insights into how commensal bacteria defend against pathogenic invasion.

**Figure 1:**
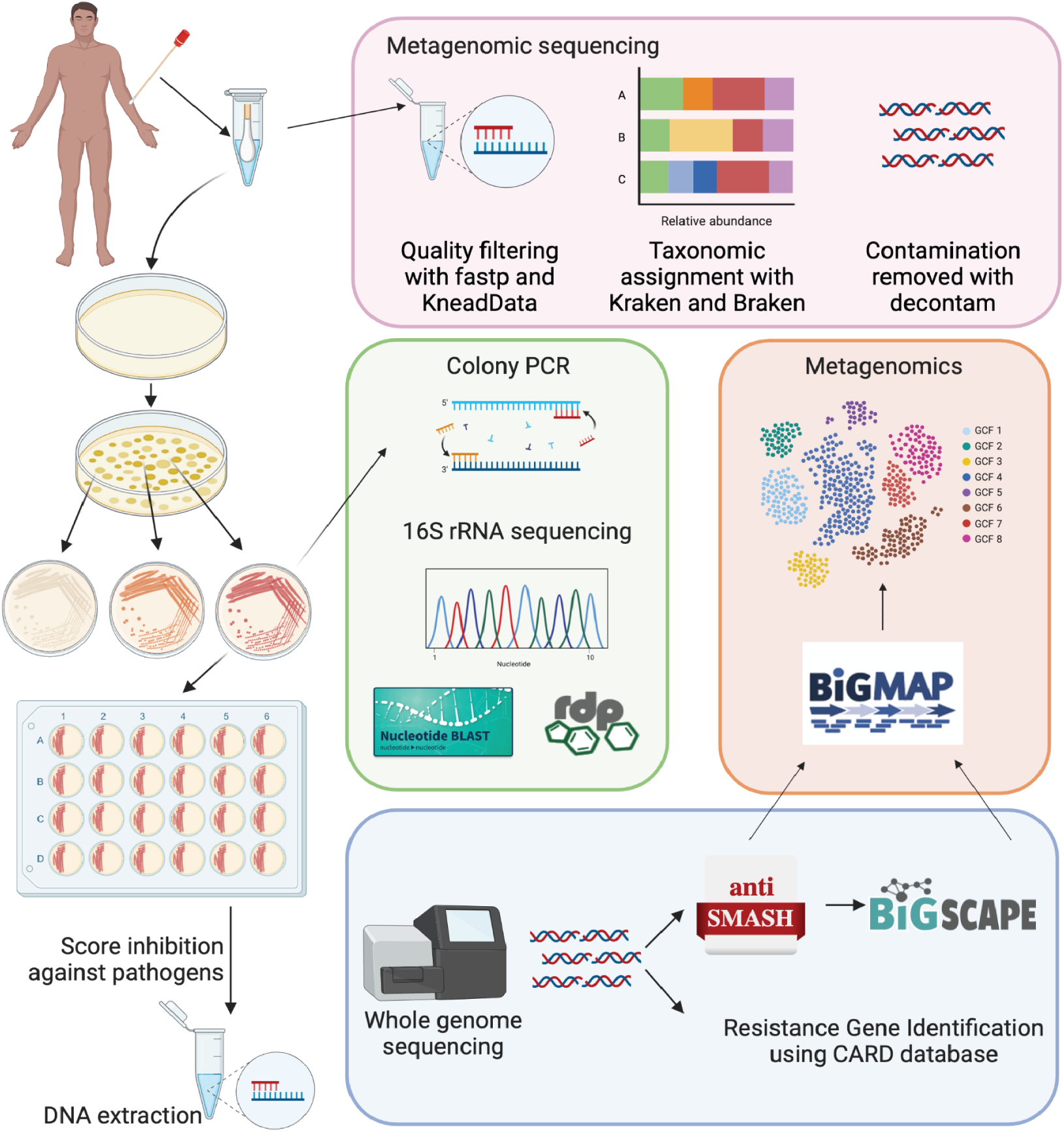
Comprehensive workflow for skin microbiome sampling and genomic analysis. Skin samples were collected from each participant from eight different body sites for metagenomic sequencing. A second set of samples was collected for strain isolation. Metagenomic sequencing was performed with taxonomic assignment using Kraken2 and Bracken. For strain isolation, samples were plated on selective media to isolate pure bacterial colonies and identified by 16S rRNA gene sequencing or whole genome sequencing. Skin isolates were assessed for antimicrobial activity using a large-scale solid-phase biological assay. Biosynthetic gene clusters and gene cluster families were annotated with antiSMASH/BiG-SCAPE and BiG-MAP for genomes and metagenomes, respectively. Antibiotic resistance genes were called using the Resistance Gene Identifier (RGI) and the Comprehensive Antibiotic Resistance Gene (CARD) database.

## Results

To create our skin microbiome collection, we recruited 34 healthy volunteers to provide samples from 8 distinct body sites (**Figure 2A**) with biogeographic ranges from moist (nares, umbilicus, toe web space), rarely moist or dry (antecubital fossa and volar forearm), and sebaceous (alar crease, back, and occiput) microenvironments (**Table S1**). Samples from 17 participants were processed for culture on several media types to capture phylogenetic and strain diversity across body sites. Shotgun metagenomic sequencing from all 34 participants was used to assess microbial community composition, diversity, and agreement between metagenomes and cultured isolates from each sample. The strain repository is named the EPithelial Isolate Collection (EPIC) and contains 980 skin-associated bacterial strains from 136 skin samples. This makes EPIC much larger and more diverse than published studies^31^. For example, 74.3% of the isolates are classified outside of the skin-dominant genera *Staphylococcus* and *Corynebacterium*, compared to 23.1% of isolates in the Skin Bacterial Culture Collection (SBCC)^31^. Furthermore, EPIC contains skin isolates belonging to an additional 24 genera not represented in the SBCC (**Table S2**).

**Figure 2:**
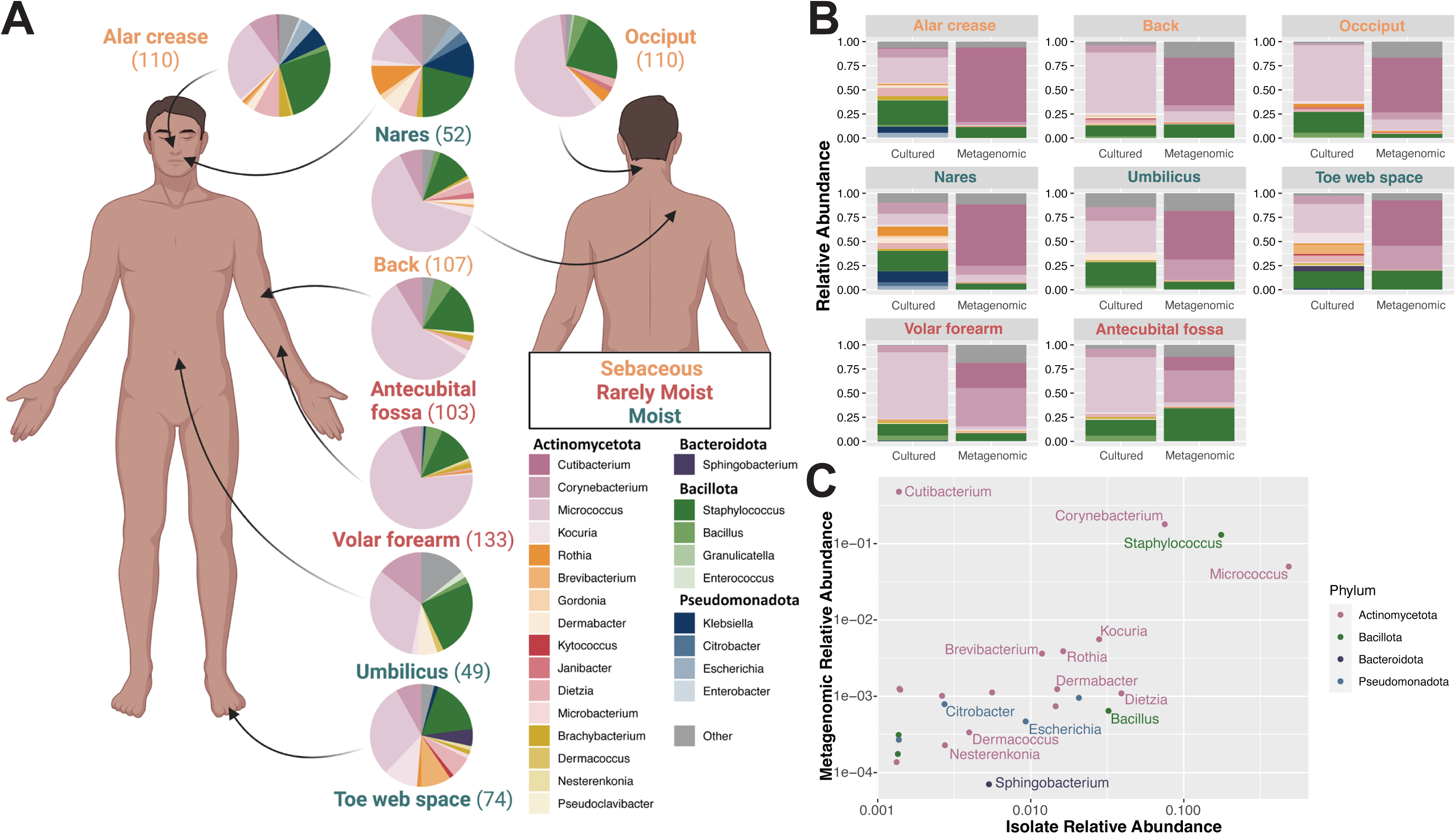
Strain library identification at each body site. (**A**) Individual pie charts represent the community structure at each site. Pie labels indicate body site, font colors indicate body site type (sebaceous, moist, rarely moist), and the n-value of each pie chart is shown in parentheses. Bacterial composition is shown within the pie charts. (**B**) Relative abundance of taxa across eight body sites comparing cultured versus metagenomic datasets. (**C**) Scatter plot depicts relative abundance of individual genera as derived from metagenomic (y-axis) and cultured datasets (x-axis). Each dot represents a specific genus, with its position reflecting the relative abundance across the two dataset types. Genera are color-coded according to their respective phyla.

### The EPIC library includes rare, low-abundance, and phylogenetically diverse bacteria

Using culture-based techniques, representative bacterial strains from each sampled body site and across individuals were isolated. To maximize the phylogenetic diversity of EPIC, each skin swab was cultured using multiple types of media, including media components to suppress *Staphylococcus* growth and to foster *Corynebacterium* and lower abundance *Actinomycetota* growth. Taxonomic classification of isolates was accomplished with full length 16S rRNA sequencing and, for a subset, whole-genome sequencing (**Table S2, S3**). In total, the 980 skin isolates represent at least 70 species (95% ANI threshold for species delineation^35^) spanning 40 genera. Most isolates fall within the phylum Actinomycetota (n=529), followed by Bacillota (n=177), Pseudomonadota (n=27), and Bacteroidota (n=4). *Micrococcus, Staphylococcus,* and *Corynebacterium* species were most frequently isolated from most body sites. We also isolated rare and phylogenetic diverse species across different genera, such as *Kocuria, Aestuariimicrobium, Kyotococcus, Nesterenkonia, Microbacterium, Brachybacterium, Rothia, Dietzia,* and *Dermabacter*. Distinct compositions of bacterial isolates were derived from each site, with the nares showing the greatest taxonomic diversity (**Figure 2A**); where 52 isolates belonging to 12 different genera were collected.

Metagenomic sequencing assessed concordance between culture-dependent versus culture-independent profiling (**Figure 2BC**). Similar to previous findings^16,36,37^, *Cutibacterium, Corynebacterium, Micrococcus*, and *Staphylococcus* are the most abundant skin colonizers across all sampled body sites. Focusing on low-abundance genera, defined as a relative abundance of < 1% in metagenomes, EPIC isolates exhibit considerable overlap with this group (**Figure 2C**). This indicates that EPIC consists of common (high abundance) skin colonizers and rare (low abundance) skin colonizers, spanning the phylogenetic diversity of the skin metagenome. Because purely aerobic culturing conditions were used, EPIC is deficient in *Cutibacterium* species due to their anaerobic nature (**Figure 2C**).

In summary, we have created a comprehensive library of skin bacterial isolates encompassing species typically present at very low levels for further exploration. We begin this exploration by: 1) mining EPIC for antimicrobial activity, using a large pairwise solid-phase screen; and 2) annotating and analyzing newly discovered BGCs.

### A pairwise interaction screen reveals widespread inhibition of pathogens by skin-associated bacterial strains

Previous studies that investigated the bioactive and antimicrobial capacity of skin microbes have focused on highly abundant species, such as *Staphylococcus* spp. Here, we developed a pairwise interaction screen to score the ability of phylogenetically diverse taxa, including low-abundance species, to provide colonization resistance to diverse human pathogens in a contact-independent manner (**Figure 3A; Table S4**). Through co-culture on solid media, we scored pairwise interactions between all 398 isolates with a pathogen panel comprising 22 Gram-positive and Gram-negative bacteria, and fungi (8,756 total interactions). Briefly, EPIC isolates are grown for 7 days to allow accumulation of secreted metabolites, followed by inoculation with the pathogen in the same well but in a spatially distant location. Interactions are scored as neutral or no inhibition (0), partial inhibition (1), or complete or full inhibition (2) (**Figure 3A**). Widespread inhibition of Gram-positive and fungal pathogens by EPIC isolates occurred, while antagonism towards Gram-negative pathogens was less common (**Figure 3BC**). Hierarchical clustering shows that isolates not exhibiting any inhibition include diverse genera. This highlights the species- and even strain-level variation in interaction patterns, where some strains of the same species may strongly inhibit the growth of specific pathogens while others do not (**Figure 3B**).

**Figure 3:**
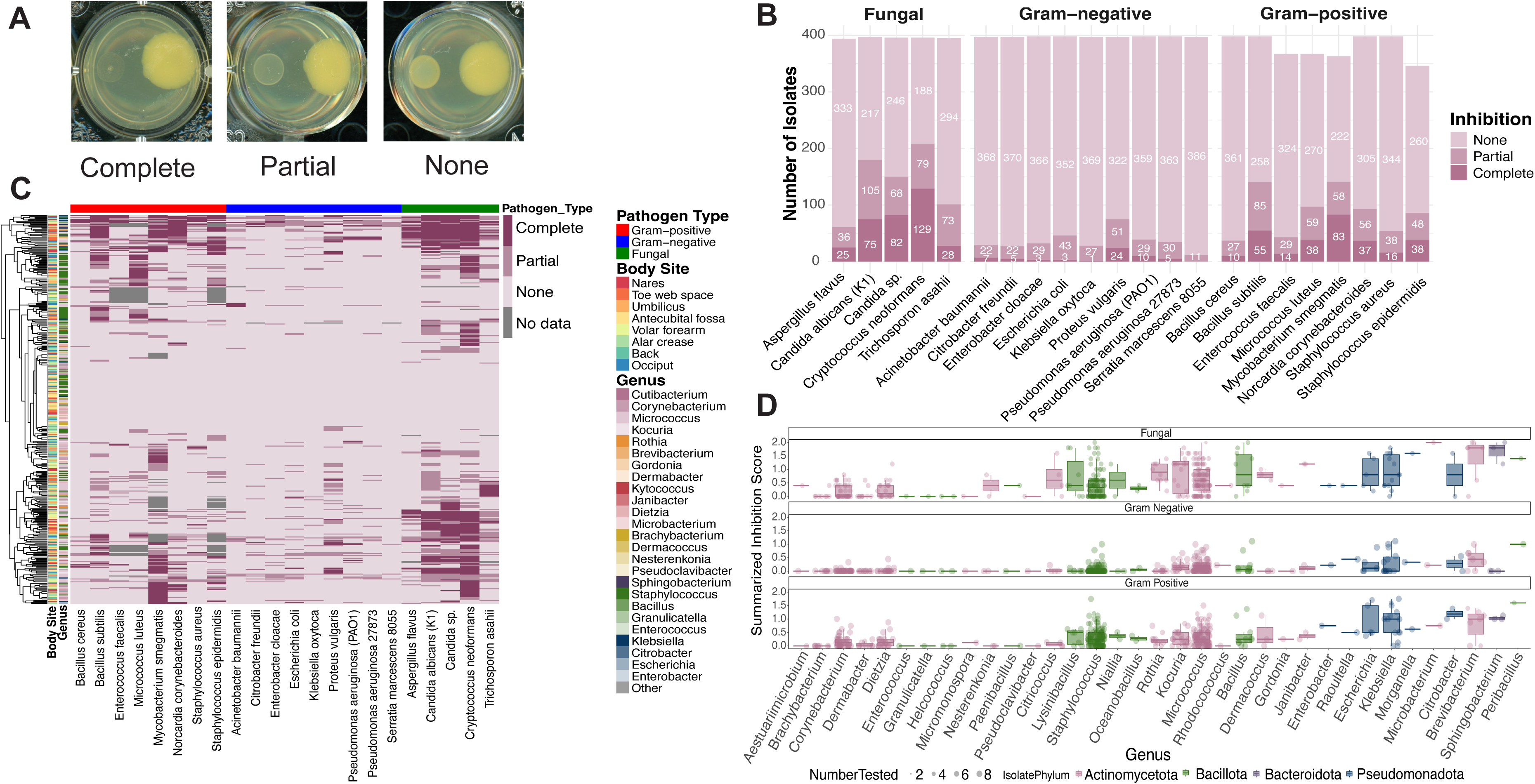
Bacteria from the human skin exhibit specific and broad antifungal activity. (**A**) Examples for each inhibition profiles: complete, partial, and none. (**B**) Quantification of inhibition (none [light pink], partial, and complete [dark pink]) profiles. Y-axis shows the number of isolates and x-axis lists members of the pathogen panel. (**C**) Hierarchical clustering of pairwise interactions where skin bacteria are tested against a panel of pathogens. Growth patterns of pathogens are scored from 0 (no inhibition) to 2 (complete or full inhibition). Rows represent individual isolates annotated by body site and genus. Columns represent a single pathogen grouped by Gram-positive, Gram-negative, and fungal species. (**D**) Inhibition score summary for each represented genus against fungal (top), Gram-negative (middle), and Gram-positive (bottom) pathogens. Dot size indicates number of isolates. Color indicates phylum of isolates.

Given the prominence of broad-spectrum fungal inhibition, we quantified the spectrum of activity for isolates inhibiting any one of the fungal pathogens. *Cryptococcus neoformans* is the most susceptible, with 129 isolates capable of completely inhibiting its growth. We identified 75 and 82 isolates able to completely inhibit the growth *Candida albicans* and *Candida* sp., respectively. Over 25 isolates have full inhibition against *Aspergillus flavus* and *Trichosporon asahii* (**Figure3B**). Together, more than 30 isolates displayed broad spectrum activity against *C. neoformans*, *Candida* sp., and *C. albicans* (**FigureS1**). A subset of 84 isolates with strong *C. albicans* inhibitory activity was randomly selected to evaluate their ability to inhibit the multidrug-resistant pathogen *Candida auris*. *C. auris* primarily spreads through skin colonization and invasive infections are associated with mortality rates exceeding 60%^38–41^. Of this subset, 40% of the isolates spanning diverse genera displayed partial to complete growth inhibition of *C. auris* (**Table S5**). Global bioactivity patterns were then assessed to determine that inhibitory profiles of EPIC isolates cluster by genus rather than body site (**Figure 3BD**). This supports a framework where closely related species display similar bioactivity profiles, independent of the local microenvironment. We developed a summarized inhibition score to further quantify antagonistic activity of each genus against each pathogen group. Isolates in the genera *Citricoccus*, *Staphylococcus*, *Kocuria*, *Micrococcus*, *Microbacterium*, *Brevibacterium*, and *Sphingobacterium* displayed the greatest fungal inhibition compared to Gram-positive and Gram-negative pathogens (**Figure 3D**). Six isolates from the genera *Brevibacterium*, *Microbacterium*, *Sphingobacterium*, and *Staphylococcus* show inhibition against all five fungal pathogens tested (**Figure S1**).

### The EPIC expands the known biosynthetic potential of the skin microbiome

Our discovery of widespread antimicrobial action among the species in our collection suggests significant novel biosynthetic capacity within the skin microbiome. To explore this biosynthetic potential, we sequenced the genomes of 287 strains displaying potent inhibitory activity. Genomes were dereplicated at 99% average nucleotide identity, resulting in a set of 182 distinct genomes for annotation of BGCs for^42^. The distribution of broad-level BGC types was similar across body sites and within taxonomic groups (**Figure 4AB**). To comprehensively understand the diversity of BGCs in skin-associated microbes, we similarly predicted BGCs across 621 genomes or MAGs from the SMGC, a recently established reference collection of microbial genomes from human skin^31^. Using unified clustering of BGCs from the SMGC and whole genomes generated in this study, we find 1,960 distinct gene-cluster families (GCFs)^43^ (**Table S6, S7**). Of the 305 GCFs found within EPIC skin isolate genomes, only 12 GCFs (3.9%) correspond to characterized BGCs from the MIBiG database, a repository of BGCs for which metabolic information is available (**Figure 4C; Table S8**). The 12 GCFs found in MIBiG, include BGCs encoded by well-studied genera associated with skin, such as BGCs for the synthesis of aureusimines^44^ and dehydroxynocardamine^45^ from *Staphylococcus* and *Corynebacterium*, respectively. The remaining known GCFs are associated with environmental taxa, such as *Bacillus* and *Streptomyces*. While some of the 305 GCFs found in EPIC genomes are also present in the SMGC, the majority, 54.4%, spanning at least 30 types of BGCs from 28 genera, are unique to EPIC (**Figure 4C**).

**Figure 4:**
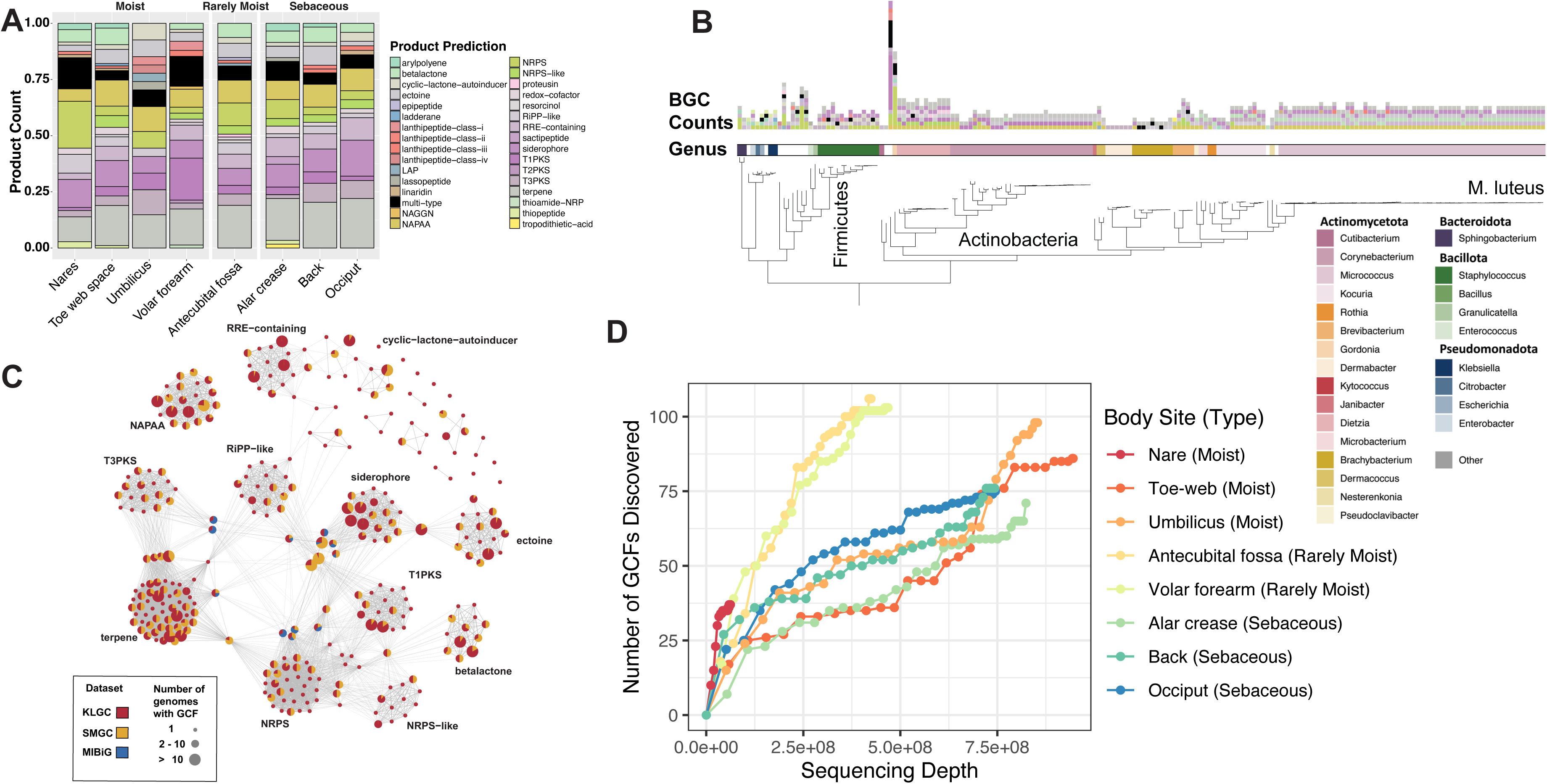
Charting biosynthetic potential of the EPIC library. (**A**) Relative abundance of different biosynthetic gene cluster (BGC) types found across eight body sites. Y-axis depicts relative abundance, while x-axis indicates distinct body sites. Each color within the bars corresponds to a unique BGC type. (**B**) A phylogeny of 182 depreplicated isolate genomes with the inner track corresponding to the genus and the outer barplot depicting the number of BGCs identified in the genome by antiSMASH. (**C**) A network of 305 GCF (nodes) identified in the genomes using BiG-SCAPE. Edges indicate common antiSMASH-based BGC types shared between GCFs. The size of the nodes corresponds to the number of genomes a GCF was found in and the color composition indicates proportion of BGCs belonging to a GCF originating from EPIC, SMGC, or MIBiG. (**D**) Cumulative GCF discovery by BiG-MAP is plotted as a function of aggregate sequencing depth at each body site across multiple individuals using skin metagenomes from Swaney et al. 2022. Metagenomes are ordered from lowest to highest sequencing depth.

To infer potential novelty of antimicrobial small molecules that may be encoded by these GCFs, we investigated the resistome of the skin microbiome, focusing on genes encoding specific antibiotic resistance enzymes that may confer self-protection^46^. Resistance to macrolides and beta-lactams were the most common across body sites; however, overall, a low prevalence of antimicrobial resistance (AMR) genes within the skin metagenome is observed (**Figure S2A**).

Similarly, through assessing whole genomes of cultured isolates, only 41 (22.5%) of 182 genomes encoded antibiotic resistance protein homologs. When considering resistance protein homologs and protein variants predicted to confer resistance, such as mutations in ribosomal protein rpsL conferring resistance to aminoglycosides, 122 of 182 genomes contained at least one resistance determinant (**Figure S2B, Table S9**).

To understand whether particular body sites exhibit greater biosynthetic potential relative to others, we examined the presence of predicted GCFs in our skin metagenomes (**Figure S3, S4**)^47,48^. The number of distinct GCFs discovered as a function of sequencing depth at each body site across multiple individuals was assessed through rarefaction^49,50^ (**Figure 4D**). Sebaceous body sites exhibit the clearest saturation for discovery of GCFs, consistent with prior observations that these sites are dominated by *Cutibacterium acnes*, which carry a limited set of GCFs^8,19,37,51^.

### Discovery of novel skin bacterial species with strong antifungal activity

Using whole genome sequencing from EPIC isolates, three *Cornynebacterium* species and one *Brachybacterium* species lacking cultured representatives but are predicted as novel species based on metagenome assembly in the SMGC were identified. In addition, four predicted novel species that are not represented in either the Genome Taxonomy Database or the SMGC were discovered. Each of these species belongs to a distinct but known genus: *Aestuariimicrobium*, *Corynebacterium*, *Kocuria*, and *Brevibacterium* (**Table S3**). To further assess the uniqueness of these species, we searched public metagenomes in the NCBI Sequence Read Archive to determine the environmental distribution^52,53^. *Corynebacterium* isolate LK952 is most commonly found in skin metagenomes, followed by gut and other human-associated metagenomes. Similarly, *Aestuariimicrobium* isolate LK1188 and *Brevibacterium* isolate LK1337 are detected in human skin metagenomes. The *Kocuria* isolate LK960 was identified in human skin metagenomes and other diverse environments, suggesting a larger host range (**Figure 5A)**.

**Figure 5:**
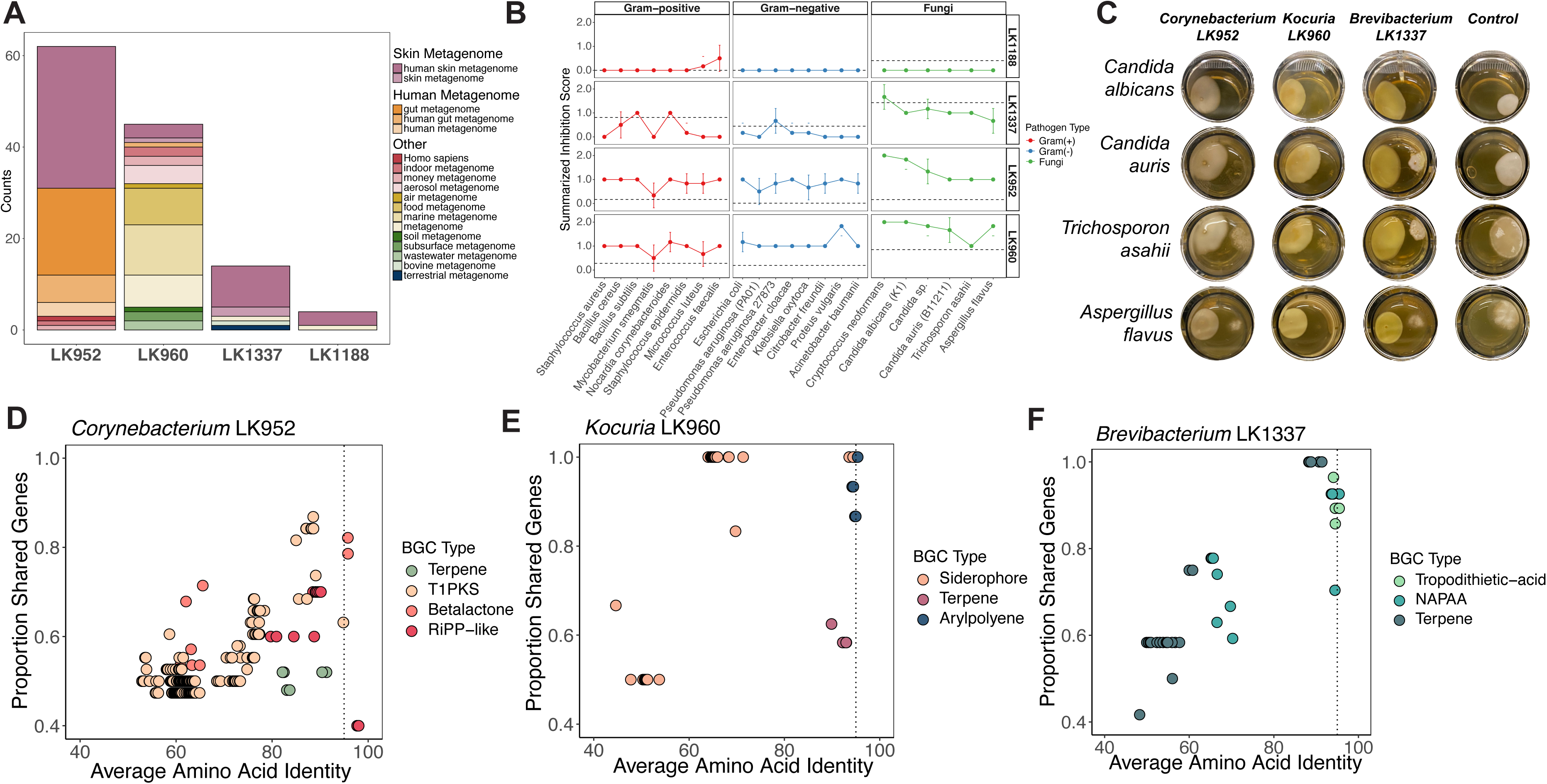
Identification of novel skin species. (**A**) Branchwater analysis of NCBI SRA metagenomes with query genomes of novel species show associations with human skin metagenomes. Colors indicate metagenome type each genome was found in. (**B**) Biological activity of *Aestuariimicrobium* LK1188 (first), *Brevibacterium* LK1337 (second), *Corynebacterium* LK952 (third), and *Kocuria* LK960 (fourth) using co-culture screening. Solid line indicates inhibition scores of against Gram-positive (red), Gram-negative (blue) and fungal pathogens (green). Dotted line indicates the average inhibition score of all isolates tested in each corresponding genus against each pathogen type. (**C**) Pairwise interaction between (in columns) LK952 (first), LK960 (second), LK13337 (third), and pathogen control (fourth) against fungal pathogens (in rows) *C. albicans* (first), *C. auris* (second), *T. asahii* (third), and *A. flavus* (fourth). (D-F) Investigating novelty of BGC-ome of (**D**) *Corynebacterium* LK952, (**E**) *Kocuria* LK960, and (**F**) *Brevibacterium* LK1337 to representative genomes from each genus. Colors indicate BGC type predicted in each species. X-axis indicates the average amino acid identity of a single BGC identified compared to the target genome. Y-axis indicates proportion of genes in identified BGC co-located in the target genome.

Each of these novel species displayed complete inhibition of fungal pathogens with limited antibacterial activity (**Figure 5B**). The *Kocuria* isolate completely inhibits *C. albicans*, *C. auris*, and *A. flavus* and although to a lesser extent, also inhibits *Trichosporon asahii* (**Figure 5C**). The novel *Corynebacterium* species exerts moderate antifungal activity while the *Brevibacterium* isolate displays less potent antifungal activity, and *Aestuariimicrobium* has no antifungal activity. Prediction and annotation of BGCs reveal the new species encode one to four BGCs each (**Figure S4**).

Finally, we assessed the novelty of each species’ BGC-ome, the collection of BGCs for a single genome, by querying the presence of predicted BGCs in genomes of known species from each genus ^54^. Intriguingly, the BGC-ome of the novel *Corynebacterium* isolate is substantially divergent from other *Corynebacterium* genomes, with several genes missing that are present within homologous BGC of other species (**Figure 5D**). *Aestuariimicrobium* contains a single BGC that is predicted to encode for a unique terpene cluster (**Table S4**). This comparative genetic analysis confirmed that BGCs encoded in each predicted novel species genomes largely have <95% amino acid identity to homologous BGCs from related species in their respective genera (**Figure 5D-F**).

## Discussion

The human skin microbiome plays an essential role in barrier maintenance and host immune regulation. In addition, it prevents pathogen invasion, indicating vast and largely unknown antimicrobial biosynthetic potential. Here, we present a human skin microbiome collection, including hundreds of strains, including cultured isolates for at least 7 new species, and metagenomes from 34 participants across 8 body sites. The isolate collection is several times larger and more diverse than existing resources and was designed to include far more taxa that are rare or low in abundance. Using this collection, we explore the expansive degree of antagonism skin microbes can exhibit towards human pathogens. Using large-scale solid-phase bioassays, we tested hundreds of strains for activity against 22 pathogens, and after assessing 8,756 pairwise interaction, we discover broad antimicrobial activity, most notably antifungal activity, in scores of strains that include new species. This confirms extensive antimicrobial biosynthetic potential, which we then explore in individual genomes, describing numerous new BGCs across multiple genera. Our collection thus has significant potential for the expansion of our knowledge of the biosynthetic resources within the skin microbiome.

Recently, broad intraspecies antagonism on the skin was reported within *S. epidermidis* strains isolated from facial skin^55^. Our findings reveal broader antagonistic networks across taxonomic lineages. Skin isolates spanning four bacterial phyla illustrate extensive cross-kingdom antimicrobial capacity with enrichment against fungal pathogens. This further suggest that inter-kingdom antagonism may be as significant as intraspecies competition in shaping microbial community assemblages on human skin.

Our resource is one of the most diverse human barrier site isolate collections. Nevertheless, *Micrococcus* is overrepresented in our strain collection. This finding is consistent with previous studies in isolating *Micrococcus* from human skin^56,57^. They are prevalent members of the skin microbiome, strict aerobes, and non-fastidious. *Micrococcus* are frequently found in indoor air^58^, which could be the result of skin shedding. Despite the overrepresentation, our culture isolation methods also captured fastidious bacteria, including novel, rare, and low-abundance skin species that appear at less than 1% abundance in metagenomes.

Based on whole-genome based taxonomic classification, we determined that eleven isolates are representative of eight novel species belonging to distinct but known genera including *Aestuariimicrobium*, *Corynebacterium*, *Kocuria*, *Brevibacterium,* and *Brachybacterium*. Six of these isolates, representing 4 novel species genomes, lacked culture representatives. Five additional isolates representing another 4 novel species, lacked both culture representatives and genomes. Three of the species in the genera *Corynebacterium*, *Kocuria*, and *Brevibacterium,* exert partial to complete inhibition of human fungal pathogens, while exerting no inhibition of Gram-positive and Gram-negative pathogens. We find that BGCs from all four novel species exhibit both substantial amino acid sequence divergence and gene content variation relative to orthologous BGCs in other species from their genera. This is likely due to genetic drift or other evolutionary processes that occur during the process of speciation ^59^. Thus, compared to the extensively studied filamentous actinomycetes, which have been heavily mined ^60^, the BGCs from rare and low-abundant genera on the skin have received less attention but are likely to be valuable sources of novel antimicrobials^61^. Our finding demonstrates that the skin microbiome can serve as a rich reservoir for exploring such rare actinomycetes and our collection makes this possible.

By surveying BGC distributions of isolates across 8 body sites, we found that the skin microbiome encodes broad classes of BGCs. This finding expands our knowledge of the distribution of BGCs in the human microbiome, including gut, vagina, airways, skin, and oral^17^. At least 32 BGC types and 305 distinct GCFs were identified within genomes of sequenced skin isolates, spanning body sites and microenvironments, suggesting the presence of a rich repertoire of specialized metabolites. Further, this indicates that there is a high degree of specialization among the microbial inhabitants, shaping skin microbiome composition through establishing colonization resistance. In addition, our analysis identified GCFs that belong to environmental taxa such as *Bacillus* and *Streptomyces*. This observation might be attributed to the skin’s continuous exposure to the environment, which potentially allows for the, usually transient, residence of environment-associated taxa and their respective BGCs on the skin^62^.

Most of the GCFs identified in EPIC skin genomes are distinct from previously-characterized BGCs, with 166 GCFs absent in skin-associated microbial genomes from the SMGC^31^. Through assessment of metagenomes, we find that rarely moist or dry sites had the highest numbers of GCFs discovered as a function of sequencing depth, suggesting that these sites have the greatest potential for isolating microbes with novel chemistries. Notably, our approach for profiling the presence of GCFs, using a catalog gathered from the EPIC and the SMGC, is likely to miss some BGCs from taxa not represented across the two genome collections.

Our understanding of the antibiotic resistome offers a strategic approach to natural product discovery and target identification^63^. Self-resistance machinery in the producer organism typically co-localize within BGCs encoding enzymatic modules to synthesize the active compound^63^. Using self-resistance as a guide has led to the discovery of thiotentronic acid antibiotics through identification of a putative fatty acid synthase resistance gene in *Salinospora*^64^ and pyxidicycline through pentapeptide repeat proteins in *Myxobacteria*^65^. We evaluated the presence of AMR genes within our skin bacterial isolates and metagenomic samples and find that the prevalence of AMR genes within the skin metagenome is low across body sites. Overall, these findings are consistent with reports that healthy host-associated microbiomes have a lower prevalence of AMR genes compared to diseased states^66,67^. This suggests that antimicrobial molecules produced likely target alternative mechanisms from currently available antibiotics, which emphasizes the need for characterizing antimicrobial molecules and their mode of actions on the human skin.

Human skin is a critical defense barrier, hosting a unique microbiome that can produce specialized metabolites to protect their niche and in turn the host. This study revealed a large and phylogenetically diverse set bacterial species with significant antifungal activity. These findings expand our understanding of antimicrobial production within the human microbiome across skin^18,19,25,68^, oral^69^, and nasal ecosystems^45^. While the broad ecology and colonization resistance function of the skin microbiome is well documented, its specific role in defense against fungal pathogens is less understood. This is a critical gap given that fungi are implicated in numerous dermatological conditions, such as pityriasis versicolor, seborrheic dermatitis, atopic dermatitis^70,71^, and chronic wounds^72,73^, but they also exist as commensals within the skin microbiome^3^. Our findings demonstrate the remarkable capacity of skin-associated bacteria to inhibit fungal growth and expansion through production of secreted antifungal molecules. This opens new lines of investigation for identifying novel and safe antifungal compounds serving a dual role of regulating the commensal fungal communities and preventing the proliferation of pathogenic fungi on the skin^3^.

## Methods

### Participant recruitment

We recruited participants at the University of Wisconsin-Madison under an Institutional Review Board approved protocol. Inclusion criteria included age >18 yr. Participation in the study was completely voluntary and participants were able to stop at any time. Participants received no payment for being a part of the study.

### Sample collection

We collected samples for metagenomic sequencing by wetting a sterile foam swab in nuclease-free water and swabbing approximately a 1 in. x 1 in. area of the selected site on the right-hand side of the participant’s body. We swabbed the area approximately 15 times in a downward motion with constant pressure while rotating the swab. We collected the swab into a 2.0 ml BioPure Eppendorf tube containing 300 μl Lucigen MasterPure™ Yeast Cell Lysis solution. Tubes were labeled with subject ID, body site, visit number, and date of collection and stored at −80 until DNA extraction.

For culturing samples, we used the Copan Diagnostics ESwab. The swab was wet in nuclease-free water, and the sample was taken from approximately a 1 in. x 1 in. area of the selected site on the left-hand side of the participant’s body. We swabbed the area approximately 15 times in a downward motion with constant pressure while rotating the swab. We then placed the swab into the Copan Diagnostics ESwab collection tube, which contained 1 ml of liquid Amies media. We labeled the tubes with subject ID, body site, and date of collection and stored at 4 for up to 24 hours until processing.

### Strain isolation and storage

Within 24 hours of collection, we added 100 μl of the Amies media from the culture sample collection tube to 900 μl of sterile water to create a 1:10 dilution. We then transferred 100 μl of the diluted sample to each of 3 agar plates: Brain Heart Infusion (BHI) + 50 mg/L mupirocin, BHI + 0.1% Tween 80 + 50 mg/L mupirocin, and Trypticase Soy Agar with 5% Sheep’s blood (blood agar) + 50 mg/L mupirocin. Blood agar plates were purchased premade, and we spread 750 μl of 1 mg/ml mupirocin to the top of the agar and allowed to soak into the media. We distributed the diluted sample evenly across the plate using sterile glass beads. We incubated the inoculated plates at 28°C for 48 to 72 hours, depending on colony formation. After incubation, we chose distinct colonies based on color, size, morphology, and opacity and struck onto a new BHI + 0.1% Tween 80 plate. We grew these strains for 24 hours or until fully grown. If the isolate was not a pure culture, we replated the isolate until it was a pure culture. Once a pure culture was obtained, we inoculated an overnight liquid culture in 3 ml BHI + 0.1% Tween 80. We stored our strains long-term at −80°C in a 2.0 mL cryotube by combining 900 μL overnight liquid culture with 900 μL 30% glycerol.

### Strain library

Of the strains isolated in our study, 451 underwent 27F 16S rRNA gene Sanger sequencing (Functional Biosciences, Madison, WI; see below). We classified sequences to genus-level using either RDP^74^ or NCBI Blast. We identified 279 isolates with whole genomes to the species level by running their genomes through autoMLST^75^.

### Colony PCR

We performed Colony PCR on the overnight liquid culture of 451 isolates in a 25μL reaction containing 12.5μL EconoTaq® PLUS 2X PCR Master Mix by Lucigen, 1μL of 10μM 27F 16S rRNA primer, 1μL of 10μL 1492R 16s rRNA primer, 10μL nuclease-free water, and 0.5μL of the overnight liquid culture. We amplified the 16S rRNA gene using the following settings: initial denaturation at 95 for 10 minutes, followed by 30-40 cycles of 95 for 30 seconds, annealing at 54 for 30 seconds, and extension at 72 for 60 seconds, with a final extension at 72 for 5 minutes and a hold at 4 indefinitely. We confirmed amplification of the 16s rRNA gene by gel electrophoresis.

### 16S Sanger sequencing

We cleaned the PCR product using the Sigma-Aldrich GenElute PCR Clean-Up kit, following kit directions. We submitted clean PCR product to Functional Biosciences in Madison, WI for Sanger sequencing of the 27F end^76^. Quality trimmed FASTA files were inputted into RDP Classifier for genus-level identification^74^.

### Microbiome DNA extractions

We performed microbiome DNA extractions on a set of swabs collected into 300uL Lucigen Master-Pure Yeast Cell Lysis buffer and frozen at −80°C. Samples were thawed on ice prior to extraction. We incorporated the extraction methods and data from Swaney and Kalan (2022)^77^. We captured the community composition across body sites and microenvironments through metagenomics profiling.

### Bioassay

Fresh bacterial isolates were struck from isolation plates or from freezer stock. We tested the bioassays in 12-well plates containing 3mL BHI solid agar in each well. A single colony from each skin isolate was streaked in a half-moon shape onto the left half each well of two 12-well plates. We incubated the plate for 7 days at 28°C. After 7 days, we spotted 3 µL of a 1:10 dilution of overnight liquid cultures from 22 different pathogens onto the right side of the well.

Two 12-well control plates with media along were included as a positive control for pathogen growth. Plates were again incubated for 7 days before scoring inhibition of growth on a scale from 0 to 3. The scoring scale was as follows; 0 – No inhibition, 1 – Slight inhibition, or more transparent than control, 2 – Medium inhibition, or zone of inhibition, 3 – Full inhibition, no pathogen growth. Inhibition scores were further simplified, grouping slight and medium inhibition together. The simplified scores are as follows: 0 – No inhibition, 1 – Slight inhibition, 2 – Full inhibition. Photos were taken of each well and uploaded along with scores to a database.

To increase throughput, we adapted the bioassay method to using 24-well plates in the spring of 2020. Liquid overnight cultures of skin isolates are grown in 3 mL BHI + 0.1% Tween 80. The bioassays were done in 24-well plates containing 1.5 mL 0.5X BHI + 0.1% Tween solid agar in each well. We spotted 1.5µL of the liquid overnight skin isolate culture onto the left half of the well. Plates were incubated at 28°C for 5 to 7 days. Slow-growing isolates were inoculated on day 0, while fast-growing isolates were introduced on day 2. On day 7, we inoculated 1µL of 1:10 diluted overnight pathogen cultures onto the right side of each well. Pathogens include Gram-positive bacteria (*Bacillus cereus*, *Bacillus subtilis*, *Enterococcus faecalis*, *Micrococcus luteus*, *Mycobacterium smegmatis*, *Staphylococcus aureus*, *Staphylococcus epidermidis*), Gram-negative bacteria (*Acinetobacter baumannii*, *Citrobacter freundii*, *Enterobacter cloacae*, *Escherichia coli*, *Klebsiella oxytoca*, *Proteus vulgaris*, *Pseudomonas aeruginosa* PAO1, *Pseudomonas aeruginosa* 27873, *Serratia marcescens* 8055), and fungi (*Aspergillus flavus*, *Candida albicans* K1, *Candida* sp., *Cryptococcus neoformans*, *Trichosporon asahii*, and *Candida auris* B11211). Plates were incubated at 28°C for 3 days. After 3 days of incubation, skin isolates were scored as before.

### gDNA extractions for whole genome sequencing

We extracted bacterial gDNA from plated isolates using the Sigma-Aldrich GenElute Bacterial Genomic DNA Kit. Library preparation and whole-genome sequencing on Illumina NextSeq 550 were performed at the SeqCenter sequencing facility (Pittsburgh, PA, USA).

### Genome assembly, representative selection through dereplication analysis, and phylogeny construction

We processed sequencing data for quality and adapters using fastp (v0.20.0)^78^ with parameters “--detect_adapter_for_pe -f 20”. Subsequently, short-read assemblies were constructed using Unicycler (v0.4.7)^79^ with default settings. Dereplication of the full set of 287 genomic assemblies at 99% identity using dRep (v3.2.2)^80^ with parameters “--S_algorithm fastANI -sa 0.99” led to the selection of 182 distinct representative genomes (**Table S2**). Genomes were taxonomically classified using GTDB-Tk (v1.7.0)^81^ with GTDB release 202^82^. GTDB-Tk was unable to assign species designations for 11 genomes. To further assess whether these genomes corresponded to novel species, we computed their average nucleotide identity comparing to genomes in the recently established SMGC database^31^ using FastANI^35^ (**Table S3**). Six of the genomes featured >95% average nucleotide identity (ANI) and >20% coverage to genomes from the SMGC and were thus regarded as known species. Genomic dereplication further indicated that two of the five remaining genomes represented the same novel species belonging to the genus of *Kocuria.* A phylogeny of the EPIC was created based on universal ribosomal proteins with GToTree (v1.6.36)^83^ and pared down to representative isolates from dereplication using PareTree^84^. Visualization of the phylogeny was performed using iTol^85^.

### Annotation of biosynthetic gene clusters, determination of gene cluster families and subsequent profiling in metagenomes

We used antiSMASH (v6.0.0) to predict the presence of BGCs for our dereplicated set of isolate genomes as well as metagenomic assembled genomes (MAGs) from skin microbiomes by Kashaf *et al.* 2022 using the parameters: “--taxon bacteria --genefinding-tool prodigal --fullhmmer --asf --cb-general --cb-subclusers --cb-knownclsuters --cc-mibig --rre --pfam2go”. Following BGC annotation, we grouped analogous BGCs across genomes into GCFs using BiG-SCAPE(v1.1.4)^43^ with requests for inclusion of singletons, hybrid classifications to be turned off, and a mixed analysis to be performed. A network graph with GCFs as nodes was constructed through parsing BiG-SCAPE mix clustering results and linking nodes based on shared annotations for GCFs stemming from antiSMASH type classifications of member BGCs. Visualization of the network was performed using igraph with the layout layout_nicely.

BiG-MAP (version committed on March 22, 2023)^48^ was used to profile the presence of BGCs from the analysis in 268 metagenomes from Swaney *et al.* 2022. We ran BiG-MAP with default parameters except for BiG-MAP.family.py, where we requested a distance cutoff of 0.3 for BiG-SCAPE based grouping of GCFs instead of 0.2. GCFs were regarded as present in a single metagenome if their full coverage was ≥ 50%, full normalized RPKM was ≥ 0.1, core gene coverage was ≥ 75% and core gene normalized RPKM was ≥ 3.0. These parameters were selected based on manual examination of observed distributions (**Figure S4**). Using these parameters, 158 different GCFs were identified by BiG-MAP in at least one metagenome. These corresponded to 148 distinct GCFs in the original BiG-SCAPE analysis, with ten BiG-SCAPE GCFs having two representatives regarded as separate GCFs by BiG-MAP.family.py. Discordance is likely because BiG-MAP performs a preliminary collapsing of similar BGCs using MASH and runs BiG-SCAPE with only representative BGCs from this initial clustering. For rarefaction analysis of GCF discovery as a function of metagenomic sequencing depth, we referenced coarser BiG-SCAPE GCF designations of representative gene clusters measured by BiG-MAP to avoid inflated metrics of GCF discovery (**Figure 4, S5**).

### Antibiotic resistance within the skin microbiome

To predict the collection of antibiotic resistance (AMR) genes, the antibiotic resistome, of the human skin microbiome, we used the Comprehensive Antibiotic Resistance Database’s (CARD) Resistance Gene Identifier (RGI) software (v6.0.1) ^46^. Briefly, we surveyed both isolate genomes and metagenomes using RGI to assess the presence of known AMR resistance genes in the CARD database. Filtered metagenomic short read sequences were compared to the reference AMR sequences in CARD via the k-mer alignment (KMA) algorithm. Metagenomes were considered to have the AMR gene present if metagenomic reads covered greater than or equal to 70% of the reference CARD AMR sequence and MAPq score greater than or equal to 50. AMR genes were grouped by the class of drugs they conferred resistance to and if a gene confers resistance to multiple drugs it counted toward both. Prevalence for antimicrobial resistance across each body site was calculated (number of subject samples from the body site with a gene confirming resistance to an antibiotic divided by the total number of subject samples from the site [n =34]).

To assess whole bacterial isolate genomes, we searched for “Perfect” and “Strict” matches of AMR genes in the CARD. We further required the percentage length of the reference sequence and identity of matches to be ≥90%. We grouped AMR genes by the class of drugs they conferred resistance to and if a gene confers resistance to multiple drugs it counted toward both. The prevalence of antimicrobial resistance within the bacterial isolates from each species was calculated by the number of isolates, with at least one gene confirming resistance to an antibiotic, divided by the total number of species isolates assessed.

### Assessment of novel species distributions across public metagenomes and comparison of their biosynthetic content to BGCs from known species in their respective genera

For each of the four novel species identified, the Branchwater webserver (https://branchwater.sourmash.bio/; accessed October) ^52^ was used to identify public metagenomes from NCBI’s SRA database which feature them. To regard a species as present in a metagenome, we required a cANI value of at least 95% to our query genome, a threshold commonly used to delineate species^86^, and a containment value of at least 0.5. The abon program, within the zol suite (v1.3.11) ^54^, was used with default settings to assess the conservation of BGCs from each novel species across genomes for other species belonging to the same genus. Comprehensive databases were independently set up for each genus by gathering genomes belonging to them from GTDB R214^87^.

## Data availability

Supplemental Material S1 to S9 can be found on GitHub (https://github.com/Kalan-Lab/SkinBioassayStudy) as SuppTable.xlsx. Whole genome assemblies are publicly available from NCBI under BioProject PRJNA803478. Metagenomic sequencing data are publicly available in the Sequence Read Archive (SRA) under BioProject PRJNA763232.

## Code availability

Code used for analyses and figures is available on GitHub (https://github.com/Kalan-Lab/SkinBioassayStudy).

## Funding and Acknowledgements

This publication was supported by the National Institutes of Health for the Biotechnology Training Program at the University of Wisconsin-Madison (5T32GM135066) [U.T.N], the National Science Foundation Graduate Research Fellowship Program [U.T.N], the National Institutes of Health awards NIAID U19AI142720 [L.R.K] and NIGMS R35GM137828 [L.R.K.], and the Weston Family Foundation Microbiome Catalyst Program [L.R.K]. The content is solely the responsibility of the authors and does not necessarily represent the official views of the National Institutes of Health.

We gratefully acknowledge members of the Kalan laboratory for discussion and feedback.

## Notes

### Competing Interest Statement

The authors have declared no competing interest.

## References

1. Swaney, M. H. & Kalan, L. R. Living in Your Skin: Microbes, Molecules, and Mechanisms. Infect. Immun. 89, (2021).

2. Townsend, E. C. & Kalan, L. R. The dynamic balance of the skin microbiome across the lifespan. Biochem. Soc. Trans. 51, 71–86 (2023).

3. Nguyen, U. T. & Kalan, L. R. Forgotten fungi: the importance of the skin mycobiome. Curr. Opin. Microbiol. 70, 102235 (2022).

4. Schommer, N. N. & Gallo, R. L. Structure and function of the human skin microbiome. Trends Microbiol. 21, 660–668 (2013).

5. Gallo, R. L. Human Skin Is the Largest Epithelial Surface for Interaction with Microbes. J. Invest. Dermatol. 137, 1213–1214 (2017).

6. Nakatsuji, T., Cheng, J. Y. & Gallo, R. L. Mechanisms for control of skin immune function by the microbiome. Curr. Opin. Immunol. 72, 324–330 (2021).

7. Almoughrabie, S. et al. Commensal Cutibacterium acnes induce epidermal lipid synthesis important for skin barrier function. Sci Adv 9, eadg6262 (2023).

8. Byrd, A. L., Belkaid, Y. & Segre, J. A. The human skin microbiome. Nat. Rev. Microbiol. 16, 143–155 (2018).

9. Harris-Tryon, T. A. & Grice, E. A. Microbiota and maintenance of skin barrier function. Science 376, 940–945 (2022).

10. Zheng, Y. et al. Commensal Staphylococcus epidermidis contributes to skin barrier homeostasis by generating protective ceramides. Cell Host Microbe 30, 301–313.e9 (2022).

11. Kengmo Tchoupa, A., Kretschmer, D., Schittek, B. & Peschel, A. The epidermal lipid barrier in microbiome-skin interaction. Trends Microbiol. 31, 723–734 (2023).

12. Ito, Y. & Amagai, M. Dissecting skin microbiota and microenvironment for the development of therapeutic strategies. Curr. Opin. Microbiol. 74, 102311 (2023).

13. Scharschmidt, T. C. et al. A wave of regulatory T cells into neonatal skin mediates tolerance to commensal microbes. Immunity 43, 1011–1021 (2015).

14. Scharschmidt, T. C. et al. Commensal microbes and hair follicle morphogenesis coordinately drive Treg migration into neonatal skin. Cell host & microbe vol. 21 467–477.e5 (2017).

15. Naik, S. et al. Commensal-dendritic-cell interaction specifies a unique protective skin immune signature. Nature 520, 104–108 (2015).

16. Grice, E. A. & Segre, J. A. The skin microbiome. Nat. Rev. Microbiol. 9, 244–253 (2011).

17. Donia, M. S. et al. A systematic analysis of biosynthetic gene clusters in the human microbiome reveals a common family of antibiotics. Cell 158, 1402–1414 (2014).

18. Zipperer, A. et al. Human commensals producing a novel antibiotic impair pathogen colonization. Nature 535, 511–516 (2016).

19. Claesen, J. et al. A Cutibacterium acnes antibiotic modulates human skin microbiota composition in hair follicles. Sci. Transl. Med. 12, (2020).

20. Nakatsuji, T. et al. Antimicrobials from human skin commensal bacteria protect against Staphylococcus aureus and are deficient in atopic dermatitis. Sci. Transl. Med. 9, (2017).

21. Nakatsuji, T. et al. A commensal strain of Staphylococcus epidermidis protects against skin neoplasia. Sci Adv 4, eaao4502 (2018).

22. Saising, J. et al. Activity of gallidermin on Staphylococcus aureus and Staphylococcus epidermidis biofilms. Antimicrob. Agents Chemother. 56, 5804–5810 (2012).

23. Nakatsuji, T. et al. Competition between skin antimicrobial peptides and commensal bacteria in type 2 inflammation enables survival of S. aureus. Cell Rep. 42, 112494 (2023).

24. Cogen, A. L. et al. Selective antimicrobial action is provided by phenol-soluble modulins derived from Staphylococcus epidermidis, a normal resident of the skin. J. Invest. Dermatol. 130, 192–200 (2010).

25. O’Sullivan, J. N., Rea, M. C., O’Connor, P. M., Hill, C. & Ross, R. P. Human skin microbiota is a rich source of bacteriocin-producing staphylococci that kill human pathogens. FEMS Microbiol. Ecol. 95, (2019).

26. Otto, M. Phenol-soluble modulins. Int. J. Med. Microbiol. 304, 164–169 (2014).

27. Okuda, K.-I. et al. Effects of bacteriocins on methicillin-resistant Staphylococcus aureus biofilm. Antimicrob. Agents Chemother. 57, 5572–5579 (2013).

28. Li, Z. et al. Integrated human skin bacteria genome catalog reveals extensive unexplored habitat-specific microbiome diversity and function. Adv. Sci. (Weinh.) 10, e2300050 (2023).

29. Salamzade, R. et al. Evolutionary investigations of the biosynthetic diversity in the skin microbiome using lsaBGC. Microb. Genom. 9, 000988 (2023).

30. Conwill, A. et al. Anatomy promotes neutral coexistence of strains in the human skin microbiome. Cell Host Microbe 30, 171–182.e7 (2022).

31. Saheb Kashaf, S., et al. Integrating cultivation and metagenomics for a multi-kingdom view of skin microbiome diversity and functions. Nat Microbiol 7, 169–179 (2022).

32. Terlouw, B. R. et al. MIBiG 3.0: a community-driven effort to annotate experimentally validated biosynthetic gene clusters. Nucleic Acids Res. 51, D603–D610 (2023).

33. Gavriilidou, A. et al. Compendium of specialized metabolite biosynthetic diversity encoded in bacterial genomes. Nat. Microbiol. 7, 726–735 (2022).

34. Uberoi, A. et al. Commensal microbiota regulates skin barrier function and repair via signaling through the aryl hydrocarbon receptor. Cell Host Microbe 29, 1235–1248.e8 (2021).

35. Jain, C., Rodriguez-R, L. M., Phillippy, A. M., Konstantinidis, K. T. & Aluru, S. High throughput ANI analysis of 90K prokaryotic genomes reveals clear species boundaries. Nat. Commun. 9, 5114 (2018).

36. Oh, J. et al. Biogeography and individuality shape function in the human skin metagenome. Nature 514, 59–64 (2014).

37. Oh, J. et al. Temporal Stability of the Human Skin Microbiome. Cell 165, 854–866 (2016).

38. Calvo, B. et al. First report of Candida auris in America: Clinical and microbiological aspects of 18 episodes of candidemia. J. Infect. 73, 369–374 (2016).

39. Lee, W. G. et al. First three reported cases of nosocomial fungemia caused by Candida auris. J. Clin. Microbiol. 49, 3139–3142 (2011).

40. Chakrabarti, A. et al. Incidence, characteristics and outcome of ICU-acquired candidemia in India. Intensive Care Med. 41, 285–295 (2015).

41. Spivak, E. S. & Hanson, K. E. Candida auris: an Emerging Fungal Pathogen. J. Clin. Microbiol. 56, (2018).

42. Blin, K. et al. antiSMASH 6.0: improving cluster detection and comparison capabilities. Nucleic Acids Res. 49, W29–W35 (2021).

43. Navarro-Muñoz, J. C. et al. A computational framework to explore large-scale biosynthetic diversity. Nat. Chem. Biol. 16, 60–68 (2020).

44. Wilson, D. J., Shi, C., Teitelbaum, A. M., Gulick, A. M. & Aldrich, C. C. Characterization of AusA: a dimodular nonribosomal peptide synthetase responsible for the production of aureusimine pyrazinones. Biochemistry 52, 926–937 (2013).

45. Stubbendieck, R. M. et al. Competition among Nasal Bacteria Suggests a Role for Siderophore-Mediated Interactions in Shaping the Human Nasal Microbiota. Appl. Environ. Microbiol. 85, (2019).

46. Alcock, B. P. et al. CARD 2023: expanded curation, support for machine learning, and resistome prediction at the Comprehensive Antibiotic Resistance Database. Nucleic Acids Res. 51, D690–D699 (2023).

47. Swaney, M. H., Sandstrom, S. & Kalan, L. R. Cobamide sharing drives skin microbiome dynamics. bioRxiv 2020.12.02.407395 (2021) doi:10.1101/2020.12.02.407395.

48. Pascal Andreu, V., et al. BiG-MAP: an Automated Pipeline To Profile Metabolic Gene Cluster Abundance and Expression in Microbiomes. mSystems 6, e0093721 (2021).

49. Chao, A. & Jost, L. Coverage-based rarefaction and extrapolation: standardizing samples by completeness rather than size. Ecology vol. 93 2533–2547 Preprint at 10.1890/11-1952.1 (2012).

50. Loureiro, C. et al. Comparative Metagenomic Analysis of Biosynthetic Diversity across Sponge Microbiomes Highlights Metabolic Novelty, Conservation, and Diversification. mSystems 7, e0035722 (2022).

51. Salamzade, R. et al. lsaBGC provides a comprehensive framework for evolutionary analysis of biosynthetic gene clusters within focal taxa. bioRxiv 2022.04.20.488953 (2022) doi:10.1101/2022.04.20.488953.

52. Irber, L., Tessa Pierce-Ward, N. & Titus Brown, C. Sourmash Branchwater Enables Lightweight Petabyte-Scale Sequence Search. bioRxiv 2022.11.02.514947 (2022) doi:10.1101/2022.11.02.514947.

53. Leinonen, R., Sugawara, H., Shumway, M. & International Nucleotide Sequence Database Collaboration. The sequence read archive. Nucleic Acids Res. 39, D19–21 (2011).

54. Salamzade, R., et al. zol & fai: large-scale targeted detection and evolutionary investigation of gene clusters. bioRxiv (2023) doi:10.1101/2023.06.07.544063.

55. Mancuso, C. P. et al. Intraspecies warfare restricts strain coexistence in human skin microbiomes. Microbiology (2024).

56. Mohapatra, N. & Kloos, W. E. Biochemical and genetic studies of laboratory purine auxotrophic strains of Micrococcus luteus. Can. J. Microbiol. 20, 1751–1754 (1974).

57. Timm, C. M. et al. Isolation and characterization of diverse microbial representatives from the human skin microbiome. Microbiome 8, 58 (2020).

58. Kooken, J. M., Fox, K. F. & Fox, A. Characterization of Micrococcus strains isolated from indoor air. Mol. Cell. Probes 26, 1–5 (2012).

59. Lassalle, F. & Didelot, X. Bacterial microevolution and the pangenome. in The Pangenome 129–149 (Springer International Publishing, Cham, 2020).

60. Hutchings, M. I., Truman, A. W. & Wilkinson, B. Antibiotics: past, present and future. Curr. Opin. Microbiol. 51, 72–80 (2019).

61. Parra, J. et al. Antibiotics from rare actinomycetes, beyond the genus Streptomyces. Curr. Opin. Microbiol. 76, 102385 (2023).

62. Mhuireach, G. Á. et al. Temporary establishment of bacteria from indoor plant leaves and soil on human skin. Environ Microbiome 17, 61 (2022).

63. Hobson, C., Chan, A. N. & Wright, G. D. The antibiotic resistome: A guide for the discovery of natural products as antimicrobial agents. Chem. Rev. 121, 3464–3494 (2021).

64. Tang, X. et al. Identification of thiotetronic acid antibiotic biosynthetic pathways by target-directed genome mining. ACS Chem. Biol. 10, 2841–2849 (2015).

65. Panter, F., Krug, D., Baumann, S. & Müller, R. Self-resistance guided genome mining uncovers new topoisomerase inhibitors from myxobacteria. Chem. Sci. 9, 4898–4908 (2018).

66. Almeida, V. de S. M., et al. Bacterial diversity and prevalence of antibiotic resistance genes in the oral microbiome. PLoS One 15, e0239664 (2020).

67. Liu, C. et al. Species-Level Analysis of the Human Gut Microbiome Shows Antibiotic Resistance Genes Associated With Colorectal Cancer. Front. Microbiol. 12, 765291 (2021).

68. O’Sullivan, J. N. et al. Nisin J, a Novel Natural Nisin Variant, Is Produced by Staphylococcus capitis Sourced from the Human Skin Microbiota. J. Bacteriol. 202, (2020).

69. Barber, C. C. & Zhang, W. Small molecule natural products in human nasal/oral microbiota. J. Ind. Microbiol. Biotechnol. 48, (2021).

70. Han, S. H. et al. Analysis of the skin mycobiome in adult patients with atopic dermatitis. Exp. Dermatol. 27, 366–373 (2018).

71. Sparber, F. et al. The Skin Commensal Yeast Malassezia Triggers a Type 17 Response that Coordinates Anti-fungal Immunity and Exacerbates Skin Inflammation. Cell Host Microbe 25, 389–403.e6 (2019).

72. Kalan, L. et al. Redefining the Chronic-Wound Microbiome: Fungal Communities Are Prevalent, Dynamic, and Associated with Delayed Healing. MBio 7, (2016).

73. Cheong, J. Z. A. et al. Priority effects dictate community structure and alter virulence of fungal-bacterial biofilms. ISME J. 15, 2012–2027 (2021).

74. Wang, Q., Garrity, G. M., Tiedje, J. M. & Cole, J. R. Naive Bayesian classifier for rapid assignment of rRNA sequences into the new bacterial taxonomy. Appl. Environ. Microbiol. 73, 5261–5267 (2007).

75. Alanjary, M., Steinke, K. & Ziemert, N. AutoMLST: an automated web server for generating multi-locus species trees highlighting natural product potential. Nucleic Acids Res. 47, W276–W282 (2019).

76. Sanger, F., Nicklen, S. & Coulson, A. R. DNA sequencing with chain-terminating inhibitors. Proc. Natl. Acad. Sci. U. S. A. 74, 5463–5467 (1977).

77. Swaney, M. H., Sandstrom, S. & Kalan, L. R. Cobamide Sharing Is Predicted in the Human Skin Microbiome. mSystems 7, e0067722 (2022).

78. Chen, S., Zhou, Y., Chen, Y. & Gu, J. fastp: an ultra-fast all-in-one FASTQ preprocessor. Bioinformatics 34, i884–i890 (2018).

79. Wick, R. R., Judd, L. M., Gorrie, C. L. & Holt, K. E. Unicycler: Resolving bacterial genome assemblies from short and long sequencing reads. PLoS Comput. Biol. 13, e1005595 (2017).

80. Olm, M. R., Brown, C. T., Brooks, B. & Banfield, J. F. dRep: a tool for fast and accurate genomic comparisons that enables improved genome recovery from metagenomes through de-replication. ISME J. 11, 2864–2868 (2017).

81. Chaumeil, P.-A., Mussig, A. J., Hugenholtz, P. & Parks, D. H. GTDB-Tk: a toolkit to classify genomes with the Genome Taxonomy Database. Bioinformatics 36, 1925–1927 (2019).

82. Parks, D. H. et al. A complete domain-to-species taxonomy for Bacteria and Archaea. Nat. Biotechnol. 38, 1079–1086 (2020).

83. Lee, M. D. GToTree: a user-friendly workflow for phylogenomics. Bioinformatics 35, 4162–4164 (2019).

84. PareTree 1.0: Remove sequences, bootstraps, and branch lengths from your trees! http://emmahodcroft.com/PareTree.html.

85. Letunic, I. & Bork, P. Interactive Tree Of Life (iTOL) v5: an online tool for phylogenetic tree display and annotation. Nucleic Acids Res. 49, W293–W296 (2021).

86. Olm, M. R., et al. Consistent Metagenome-Derived Metrics Verify and Delineate Bacterial Species Boundaries. mSystems 5, (2020).

87. Parks, D. H. et al. GTDB: an ongoing census of bacterial and archaeal diversity through a phylogenetically consistent, rank normalized and complete genome-based taxonomy. Nucleic Acids Res. (2021) doi:10.1093/nar/gkab776.

