## Supplemental Figure 1 for "Large-scale investigation for antimicrobial activity reveals novel defensive species across the healthy skin microbiome"

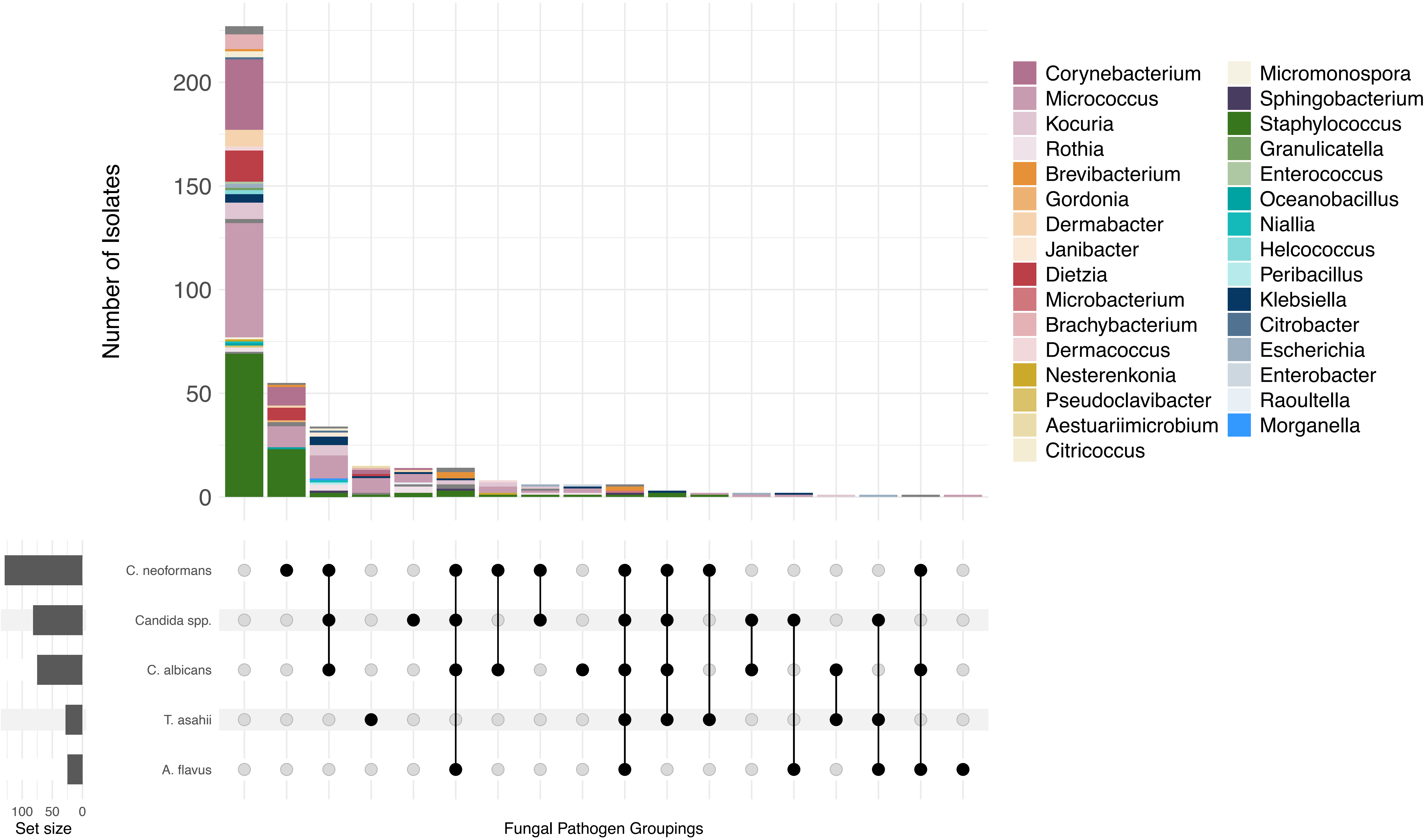


**Figure S1: Fungal inhibition by skin isolates.** The top panel illustrates a stacked bar plot, where each color corresponds to a specific bacterial genus, showing the distribution of strain counts (y-axis) within each genus. The bottom matrix focuses on the intersections among different fungal pathogen types. Rows are labelled to represent distinct fungal pathogens, and each column signifies the overlapping occurrence of these pathogens across sampled sets. Cells within the matrix are filled to illustrate the presence of a fungal pathogen type in the intersecting sets, with filled cells in the same column connected by a horizontal line. To the left of this matrix, bar charts corresponding to the row labels indicate the total number of instances for each fungal pathogen type.
