## Supplemental Figure 2 for "Large-scale investigation for antimicrobial activity reveals novel defensive species across the healthy skin microbiome"

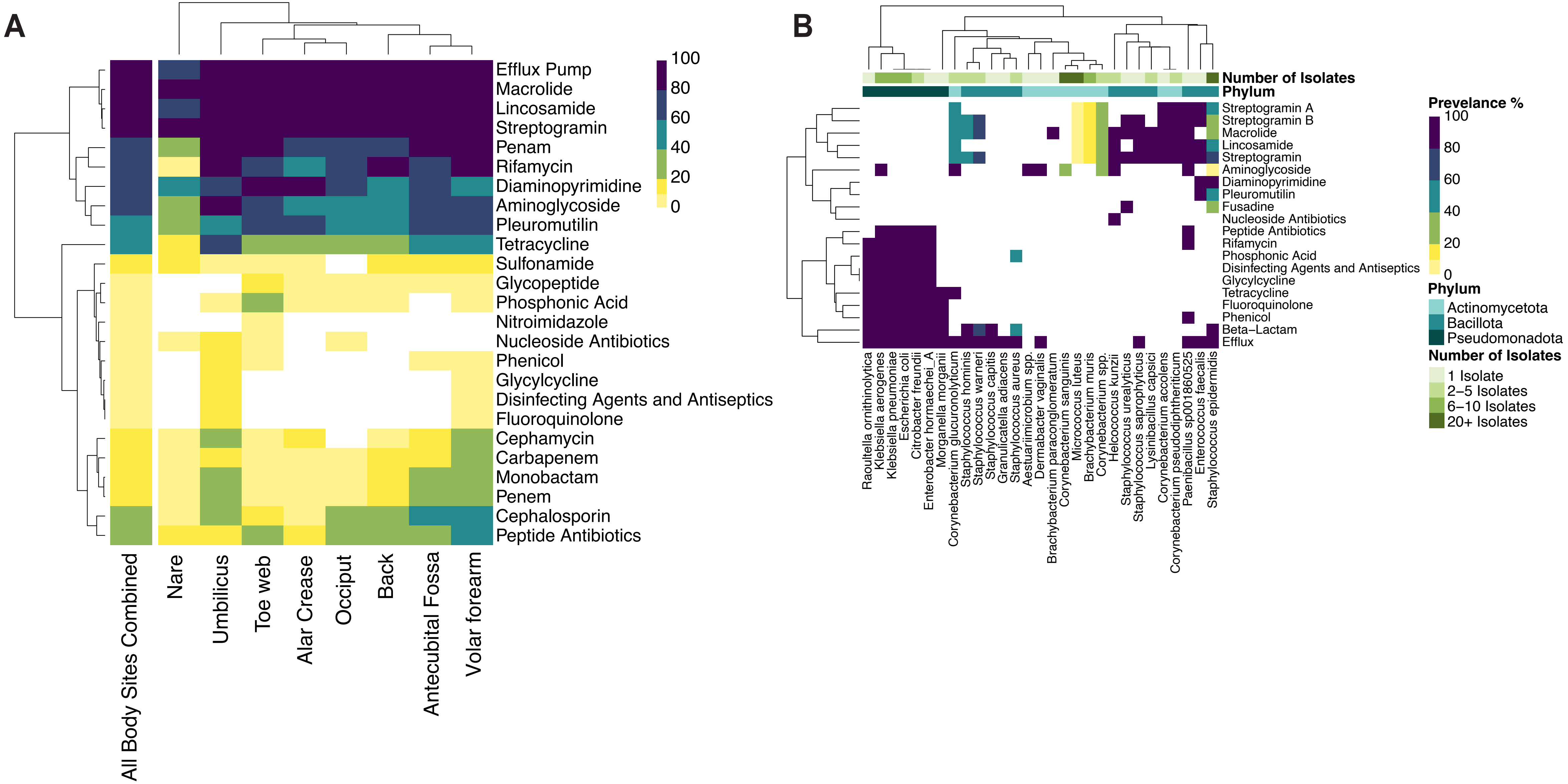


**Figure S2: Low prevalence of antibiotic resistance genes in the EPIC library.** (**A**) Prediction of antibiotic resistance genes in the skin metagenomes. Rows indicate antibiotic classes. Columns indicate body sites. Colors indicate prevalence of antibiotic resistance across each body site. (**B**) Profiling of antibiotic resistance genes in 287 whole genomes from cultured skin isolates.
