## Supplemental Figure 3 for "Large-scale investigation for antimicrobial activity reveals novel defensive species across the healthy skin microbiome"

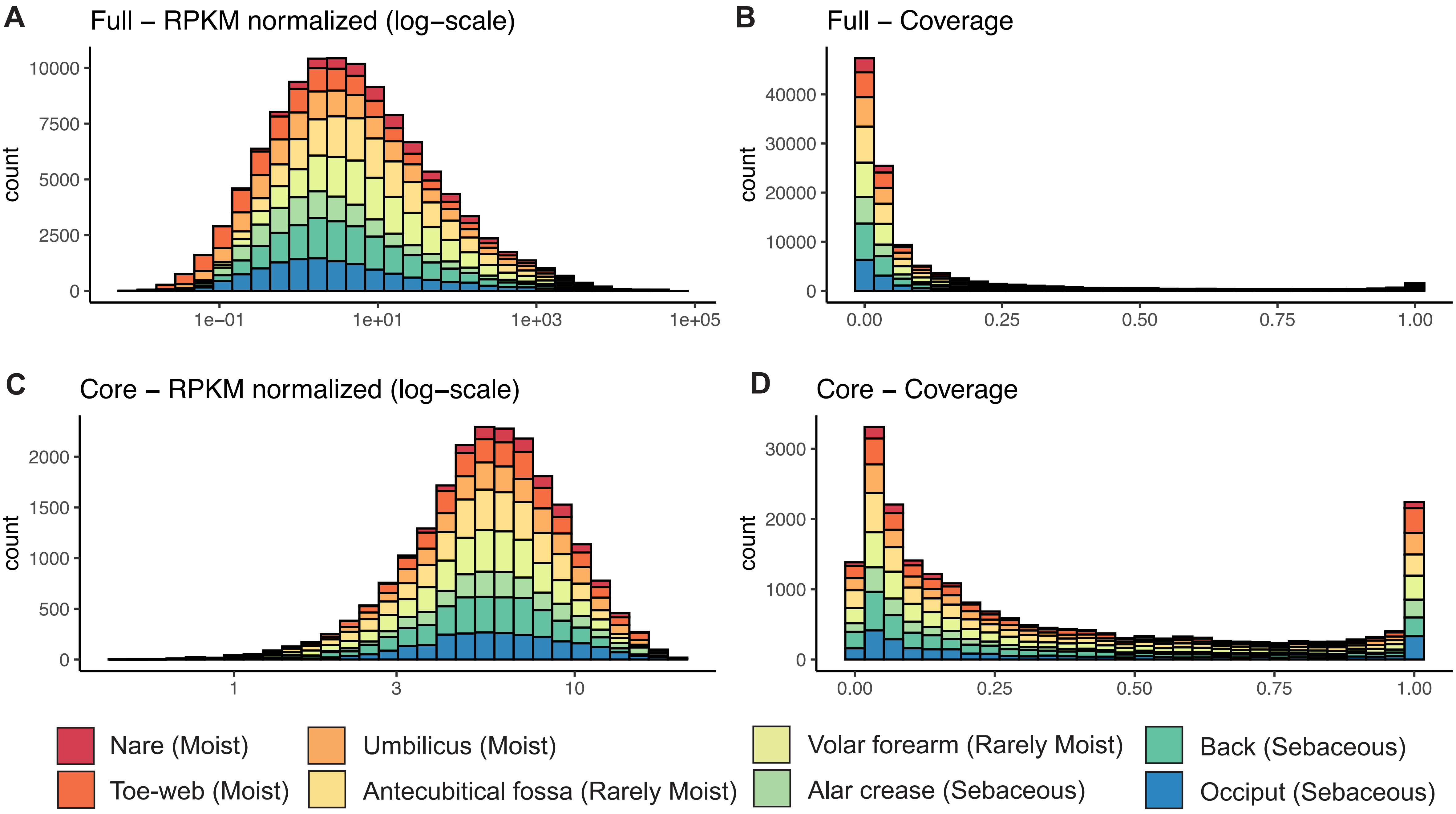


**Figure S3: Representative BGC coverage and normalized RPKM metrics reported by BiG-MAP.** Distribution of the normalized RPKM metric is shown across all metagenomes from Swaney et al. 2022 for full (**A**) and core (**B**) regions in representative BGCs. Similarly, distributions of the proportion of sites across full (**C**) and core (**D**) regions of GCFs is also shown. Only non-zero datapoints are shown.
