## Supplemental Figure 4 for "Large-scale investigation for antimicrobial activity reveals novel defensive species across the healthy skin microbiome"

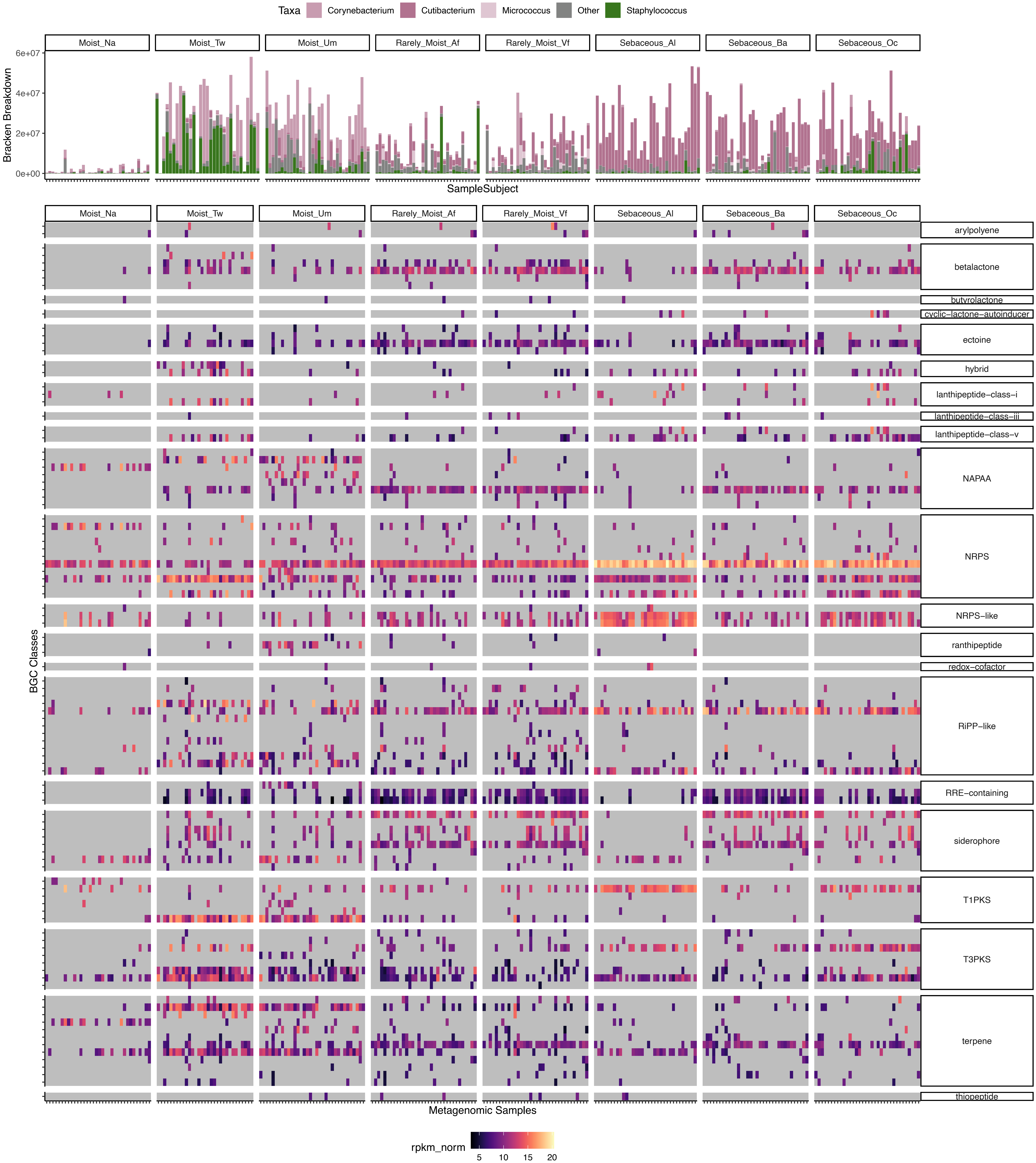


**Figure S4: The distribution of GCFs across metagenomes.** The normalized RPKM of core regions for GCFs (rows) partitioned by their BGC annotation class (row groups) is shown across metagenomes (columns) divided according to body site of sampling (column groups). Grey indicates the GCF was not detected for a particular metagenome at the required cutoffs. Only GCFs found in five or more metagenomes are shown.
