## Supplemental Figure 5 for "Large-scale investigation for antimicrobial activity reveals novel defensive species across the healthy skin microbiome"

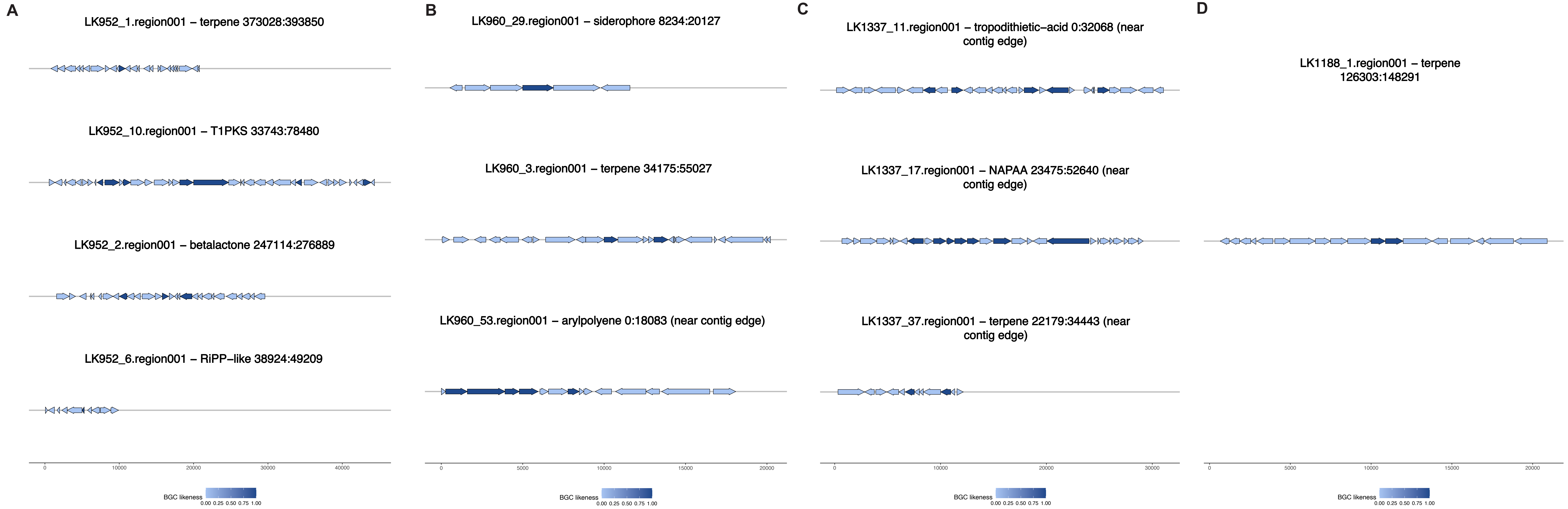


**Figure S5: Visualization of BGCs in novel, skin-associated species**, including (**A**) Corynebacterium LK952, (**B**) Kocuria LK960, (**C**) Brevibacterium LK1337, and (**D**) Aestuariimicrobium LK1188. Colors correspond to the BGC-likeliness of each gene.
